# Perceptual inference employs intrinsic alpha frequency to resolve perceptual ambiguity

**DOI:** 10.1101/399840

**Authors:** Lu Shen, Biao Han, Lihan Chen, Qi Chen

## Abstract

The brain uses its intrinsic dynamics to actively predict observed sensory inputs, especially under perceptual ambiguity. However, it remains unclear how this inference process is neurally implemented in biasing perception of ambiguous inputs towards the predicted percepts. Using electroencephalography and intracranial recordings, we first show that the alpha-band frequency defines a unified time window for perceptual grouping across both space and time: information segments, either spatially or temporally segregated, will be integrated if they fall within the same alpha cycle. Moreover, predictions employ this prior knowledge on intrinsic alpha frequency to shift perceptual inference towards the most possibly observed percepts. Multivariate decoding analysis showed that perceptual inference, based on variance in prestimulus alpha frequency (PAF), biases post-stimulus neural representations by inducing preactivation of the predicted percepts. fMRI results additionally showed that prestimulus activity and intrinsic organization status in the frontoparietal attentional network predict perceptual outcomes, probably by modulating occipitoparietal PAFs.

## Introduction

The brain is increasingly being understood as engaged in probabilistic unconscious perceptual inference to actively predict and explain observed sensory inputs, which helps, via top-down prediction signals, to resolve ambiguity in bottom-up sensory signals [1–6]. Therefore, our perception of the world is not simply based on bottom-up inputs from our sensory organs. Instead, what we perceive is heavily altered by contextual information [7–10] and expectations [11–14]. Besides context and expectation, intrinsic neural oscillatory signatures and organization status of the brain dramatically modulate the perceptual outcome of ambiguous stimuli [15–19].However, it remains unclear how perceptual inference employs intrinsic brain activity to bias the perception of ambiguous sensory information towards predicted percepts, with the progress of time.

The process of perceptual inference can be well illustrated by the phenomenon of bistable apparent motion in the Ternus display [20,21], in which subjective perception spontaneously alternates between spatial and temporal grouping interpretations of a constant ambiguous dynamic visual scene (Fig 1). The human brain adopts two major strategies of perceptual grouping to achieve perceptual coherence along the spatial and temporal dimension, despite the ever-changing visual inputs and the resulting fragmentary nature of the retinal image across space and time [22–25]. Spatially, grouping local visual elements into a holistic percept allows us to perceive scenes and objects as a whole, rather than a meaningless collection of unconnected parts [26–28]. Temporally, successive discrete visual events unfolding in time could be grouped based on temporal proximity to perceive the stability of object identity and location [24,29,30]. In the classical Ternus display (Fig 1A), two horizontally spaced discs appear at shifted locations in two successive frames. Depending on the inter-frame interval (IFI), observers typically report two distinct percepts [31,32]. Temporal grouping is explicitly dominant at short IFIs: the central overlapping discs between the two frames are temporally integrated as one single disc, and the visual persistence of the central overlapping disc makes the lateral disc in Frame 1 appear to jump across the central disc, i.e. element motion (EM, Fig 1A, upper panel; S1 Video). On the other hand, spatial grouping is explicitly dominant at long IFIs: the two discs within each frame are spatially grouped, and perceived as moving together as a group, i.e. group motion (GM, Fig 1A, lower panel; S2 Video). Most critically, when the IFI reaches a certain psychophysical threshold, the Ternus display becomes ambiguous/bistable: the report of EM (temporal grouping) vs. GM (spatial grouping) percepts randomly fluctuates on a trial-by-trial base, resulting in comparable proportions of GM and EM percepts (Fig 2A) [24,32]. Interestingly, the typical transition IFI threshold between temporal and spatial grouping in the Ternus display occurs around a time window of ∼100 ms [32,33] (see Fig 2A, B), which corresponds to the average cycle of occipital alpha-band oscillations, with peak frequencies ranging between 8 and 13 Hz (i.e. 70-120 ms cycle).

**Fig 1.**
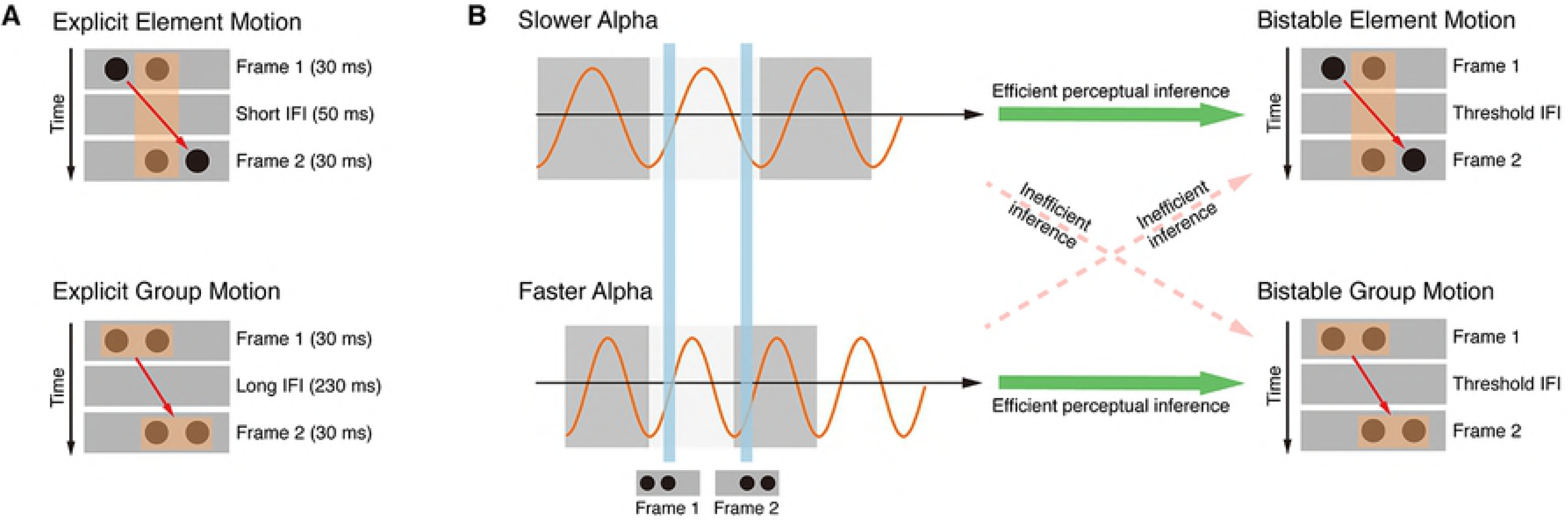
Schematic demonstration of the temporal and spatial grouping in the Ternus display and their hypothetical relations to the alpha cycles. (A) *Upper panel:* explicit element motion (EM) at the short IFI (50 ms). With the short IFI, the shared central disc between the two frames is temporally grouped as the same object, resulting in explicit EM percepts. The central disc is perceived to remain still at the central location, and the two lateral discs are perceived to jump from one side to the other (see also S1 Video). *Lower panel:* explicit group motion (GM) at the long IFI (230 ms). With the long IFI, the two discs within each of the two frames will be spatially grouped, respectively, resulting in explicit GM percepts. The two discs in the first frame are perceived to move together as a group towards the second frame in a manner consistent with the physical displacement (see also S2 Video). (B) *Upper panel:* bistable EM percepts at the threshold IFI. The lower the prestimulus alpha frequency, the higher possibility the two consecutively presented frames will fall in the same alpha cycle. Accordingly, efficient perceptual inference (green arrow) towards the EM percepts and inefficient perceptual inference (dotted red arrow) towards the GM percepts will be made. *Lower panel:* bistable GM percepts at the threshold IFI. The higher the prestimulus alpha frequency, the two frames will more likely fall in different alpha cycles. Accordingly, efficient perceptual inference (green arrow) towards the GM percepts and inefficient perceptual inference (dotted red arrow) towards the EM percepts will be made.

**Fig 2.**
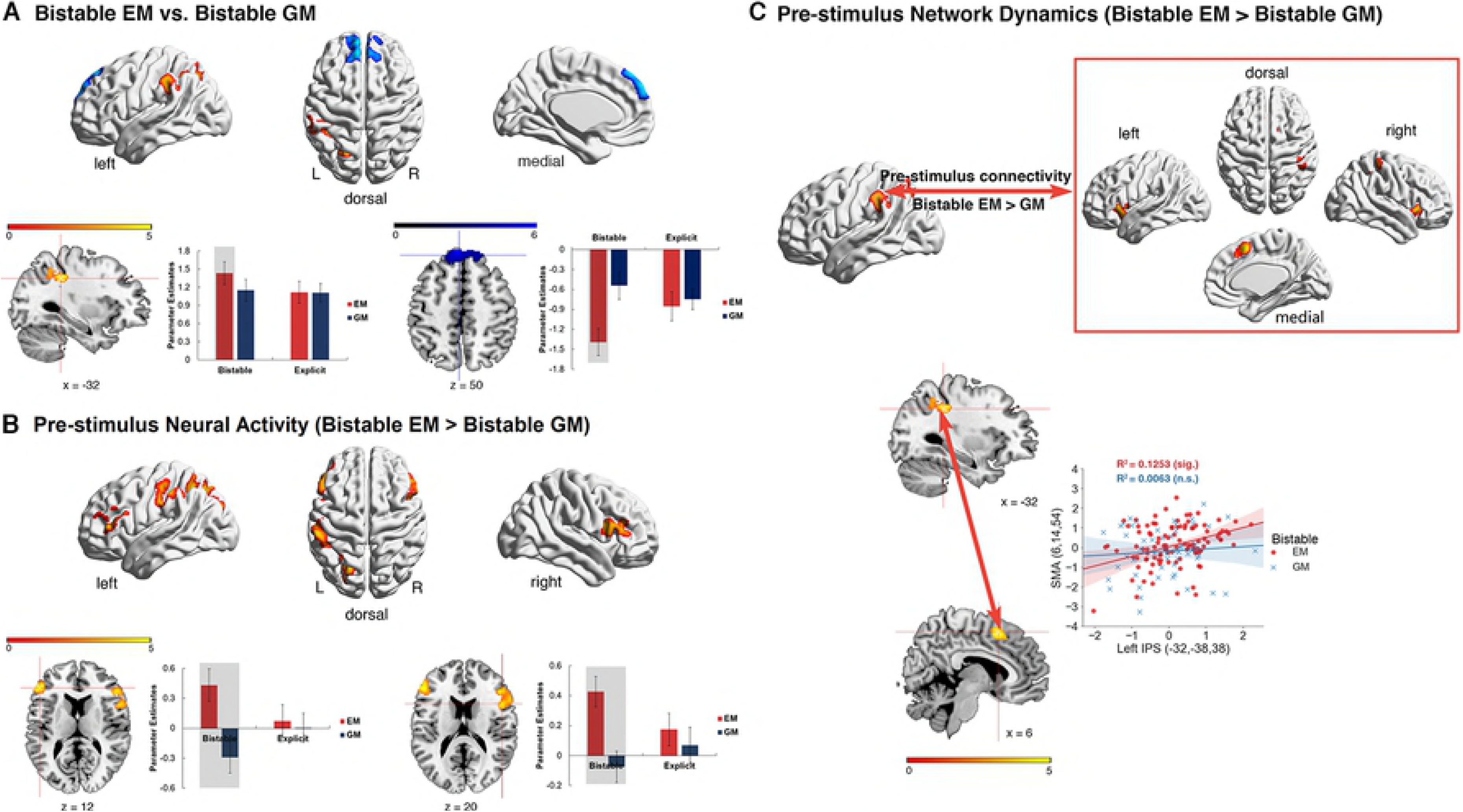
Behavioral results. (A) Psychometric curve fitted on the psychophysical data from one representative participant. The transition IFI threshold in each individual subject was defined as the IFI corresponding to the point of 50% reported GM/EM percepts on the fitted logistic function. (B) Threshold IFIs derived from the psychophysical procedure for each participant in the EEG and fMRI experiment, respectively. (C) Accuracy rates of the Explicit EM and GM trials averaged across all the participants in the EEG and fMRI experiment, respectively. (D) Rates of the Bistable EM and GM trials averaged across all the participants in the EEG and fMRI experiment, respectively. (E) Reaction times (RTs) relative to the onset of Frame 2 for all the experimental conditions in the EEG and fMRI experiment. The error bars indicate ±1 SEM. *p < 0.05, **p < 0.01.

The alpha oscillations, as one of the most predominant oscillations in the visual system, are considered as one underlying mechanism of perceptual cycles by gating the transient temporal windows of perception [34,35]. Accordingly, accumulating evidence shows that the phase of ongoing alpha oscillations reflects cyclic shifts of neuronal excitability [36–38], predicts not only behavioral performance [15,39–41], but also a variety of subsequent neural signals related to stimulus processing [16,42,43]. Besides the phasic effects, the peak frequency of alpha-band oscillations predicts reaction times [44] and variations in temporal resolution of perception [35,45–47]. The phasic and frequency effect of alpha oscillations lead to the long standing hypothesis that the alpha cycle provides the discrete temporal window of perceptual grouping: whether two stimuli are integrated into a single percept or segregated into separate events depends on whether they fall in the same cycle of the alpha oscillation [48,49]. In other words, the duration of alpha cycle provides a unified temporal window of perceptual grouping both across space and over time: any information segments (either spatially or temporally segregated) that fall in the same alpha cycle will be integrated. In terms of the Ternus paradigm, if the two frames fall in the same alpha cycle, they will be temporally integrated, resulting in the EM percepts; if the two frames fall in different alpha cycles, spatial grouping will take place separately in the two frames, resulting in the GM percepts.

Critically, when the sensory inputs become ambiguous at the transition IFI threshold, we hypothesize that perceptual inference employs intrinsic prestimulus alpha frequency (PAF) to predict the most possibly perceived percepts. Specifically speaking, lower PAFs (i.e. longer prestimulus alpha cycles, Fig 1B, upper panel) will induce the prediction that the two consecutively presented frames will more probably fall in the same alpha cycle, and accordingly perceptual inference towards the EM percepts will be made. On the other hand, higher PAFs (i.e. shorter prestimulus alpha cycles, Fig 1B, lower panel) will induce the prediction that the two frames will more probably fall in different alpha cycles, and accordingly perceptual inference towards the GM percepts will be made. We thus predict that the PAF can causally predict the outcome of bistable perceptual grouping, with higher PAFs preceding the bistable GM than EM percepts. Moreover, trial-by-trial changes in the instantaneous PAFs may induce different predictions on the corresponding spatially vs. temporally integrated percepts for perceptual inference, which may bias neural representations of the predicted percepts, with the progress of time. Especially when the PAF is obviously slow or fast, i.e. at the two opposite ends of the PAF continuum, perceptual predictions on the corresponding bistable EM (temporally integrated) vs. GM (spatially integrated) percepts will be the most efficient with the highest probabilities. Under this hypothesis, the efficient predictions may induce the corresponding representation pattern underlying the predicted percepts already before the actual presentation of the bottom-up stimuli. Alternatively, if perception was based solely on bottom-up sensory inputs, one would assume that neural representations underlying the integrated percepts are induced only after the actual presentation of the bottom-up stimuli. To distinguish between the above hypotheses, we adopted electroencephalography (EEG) in healthy adults and intracranial recordings in epileptic patients, and used multivariate decoding techniques on the EEG data to further probe the representational content of neural signals in a time-resolved manner. In addition, one functional magnetic resonance imaging (fMRI) experiment was performed to test how prestimulus intrinsic brain activity and network dynamics in the frontoparietal attentional network predict the outcome of bistable perceptual grouping.

## Results

### Behavioral performance

In the EEG (n = 17), intracranial (n = 4), and fMRI (n = 18) experiments, participants were asked to report the perceived EM vs. GM percepts after viewing (1) the explicit EM stimuli with the short IFI, (2) the explicit GM stimuli with the long IFI, and (3) the bistable stimuli with the transition IFI threshold (Fig 1; see Materials and Methods). The transition IFI threshold, at which equal proportions of EM and GM trials were reported, was determined individually for each participant prior to the main experiment (Fig 2A; see Materials and Methods). The individual IFI threshold for each participant was shown in Fig 2B for the EEG and fMRI experiments, respectively. A two-sample t-test showed no significant difference between the two experiments in terms of the group mean IFI threshold, t _(33)_ < 1.

In the two explicit conditions, the mean accuracy rate in the explicit EM and explicit GM condition was comparable, and both above 85%, in both the EEG, t _(16)_ < 1, and the fMRI experiment, t _(17)_ < 1 (Fig 2C), indicating that the participants could clearly distinguish the two explicitly different percepts at the short vs. long IFIs. Reaction times (RTs), however, were significantly slower in the explicit EM than the explicit GM condition, in both the fMRI experiment, t (17) = 3.06, p < 0.01, and the EEG experiment, t _(16)_ = 3.73, p < 0.005 (Fig 2E). In the bistable condition, there was no significant difference between the bistable EM and the bistable GM conditions, in terms of both the rates of choice, EEG experiment: t _(16)_ < 1, fMRI experiment: t _(17)_ < 1 (Fig 2D), and the RTs, EEG experiment: t _(16)_ < 1, fMRI experiment: t _(17)_ < 1 (Fig 2E). In addition, behavioral performance of the four epileptic patients with depth electrodes showed similar patterns as the healthy participants (see S1 Table).

### Neurophysiology results

#### Prestimulus alpha frequency predicted the outcome of bistable perceptual grouping

The EEG data showed a clear peak in the alpha-band amplitude (8-13 Hz) (Fig 3A), and a posterior scalp distribution of alpha amplitude during the prestimulus period (- 800 to 0 ms relative to the presentation of the first frame) for all the participants (Fig 3B). These results guided further analysis by confining the frequency of interest to 8-13 Hz, and the region of interest to the posterior electrodes (Oz, O1, O2, POz, PO1, PO2, PO3, PO4).

**Fig 3.**
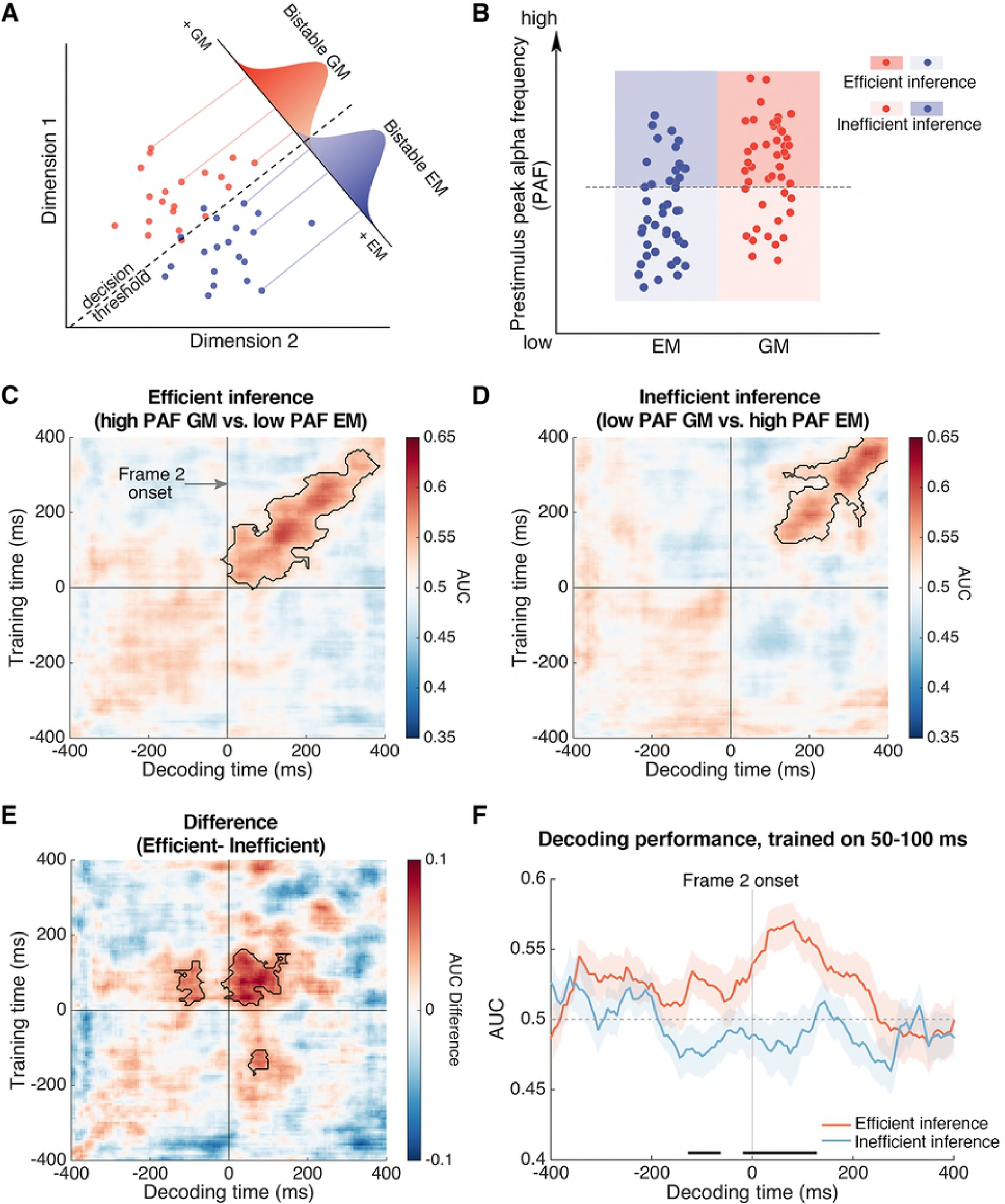
Relationship between the prestimulus alpha frequency and bistable perceptual grouping. (A) The normalized amplitude obtained through a Fast Fourier Transform (FFT) from all the bistable trials (from −800 to 0 ms relative to the onset of the first frame) collapsed over all electrodes in all the participants revealed a clear peak at the alpha frequency. The grey shading indicates ±1 SEM. The light grey rectangle indicates the selected frequency band. (B) The amplitude topographic map of the selected alpha frequency band (8–13 Hz) revealed a clear posterior scalp distribution. The selected posterior electrodes are indicated with white dots. (C) The duration of occipital alpha cycle in each individual subject, derived from the prestimulus alpha activity, was significantly correlated with the individual transition IFI threshold (rho = 0.700; p = 0.001). Dashed lines indicate 95% confidence intervals around the linear fit line. (D) Within-subject analysis of the instantaneous PAF revealed higher alpha frequency preceding the bistable GM trials than the bistable EM trials. Significant time points are indicated by the horizontal black bar (cluster-based correction, p < 0.05). Shaded regions denote ±1 within-subjects SEM.

To understand the relationship between the alpha frequency and the perceived percepts of the bistable perceptual grouping, we first calculated the between-subject correlation between the individual transition IFI thresholds and the individual prestimulus peak alpha frequency. The individual peak alpha frequency was calculated based on the maximal prestimulus posterior alpha amplitude of each participant [50]. Subsequently, the Pearson correlation between the width of individual alpha cycle and the individual transition IFI threshold was calculated. The two measures were significantly correlated, n = 17, r =.70, p =.001 (Fig 3C). The significant between-subject correlation suggests that the faster the alpha oscillations (i.e. shorter alpha cycles) in an individual, the shorter IFI is required for the two frames to fall in different alpha cycles, and thus it takes shorter IFI for the later GM percepts at the longer IFIs to take dominance over the earlier EM percepts at the shorter IFIs. On the other hand, the slower the alpha oscillations (i.e. longer alpha cycles) in an individual, the longer IFI is required for the two frames to fall in different alpha cycles, and thus it takes longer IFI for the transition from the earlier EM percepts to the later GM percepts.

Moreover, if as we predicted, slower alpha cycles lead to higher possibilities that the two frames will fall in the same alpha cycle (i.e. temporal grouping, Fig 1B, upper panel) while faster alpha cycles lead to higher possibilities that the two frames will fall in different alpha cycles (i.e. spatial grouping, Fig 1B, lower panel), trial-by-trial variance in the PAF within each subject should predict the outcome of the bistable perceptual grouping, with higher PAFs on the bistable GM (spatial grouping) than the bistable EM (temporal grouping) trials. To directly test the causal relation between the PAF and the outcome of bistable perceptual grouping, we analyzed time-resolved changes in the prestimulus derivative of the phase angle time series (see Materials and Methods), which corresponds to the instantaneous frequency of a signal within a band-limited range [51]. For each subject, we calculated and compared the instantaneous alpha frequency in the prestimulus window (−800 to 0 ms relative to the presentation of the first frame) for the bistable GM and the bistable EM percept trials, respectively. Consistent with our predictions, the results showed that the PAF was significantly higher in the bistable GM than bistable EM trials, from about −550 to −210 ms relative to the presentation of the first frame, cluster-based correction, p < 0.05 (Fig 3D).

To further test the consistency of the above PAF effect and its precise anatomical origins, we collected intracranial data from four epileptic patients with depth electrodes. For each patient, according to the present frequency of interest at 8-13 Hz, we computed the alpha amplitude for each and every contact, and then selected the first ten contacts with the strongest alpha amplitude. The anatomical locations of these selected contacts were mostly located in the occipital and parietal regions (Fig 4), which was consistent with the posterior scalp distribution of alpha in the EEG experiment (Fig 3B).

**Fig 4.**
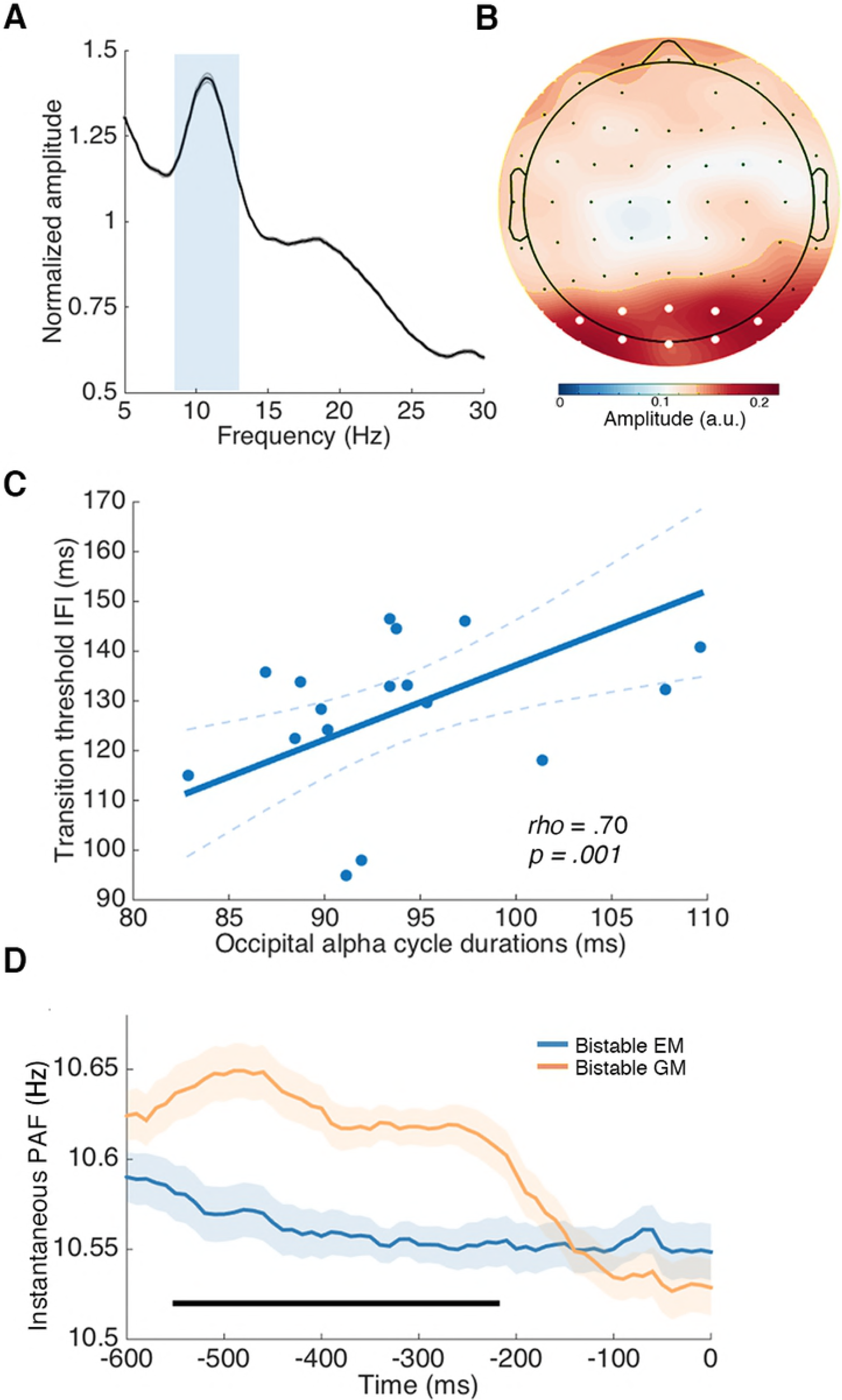
Prestimulus alpha frequency effect in the four epileptic patients. Anatomical locations of the first ten contacts with the largest alpha (8-13 Hz) power from each of four patients (A, B, C, and D). Precise anatomical regions of the ten selected contacts are marked on the coronal T1 slices of individual brains. Instantaneous PAF was calculated for the bistable EM and bistable GM trials, respectively, for each selected contact, and averaged across all the contacts. The alpha frequency preceding the bistable GM trials was higher than that preceding the bistable EM trials in most of selected contacts in each patient. In the averaged results, significant time points are indicated by horizontal black bars (cluster-based correction, p < 0.05). Shaded regions denote ±1 SEM.

We subsequently performed further analysis on the within-subject trial-by-trial variance of instantaneous PAF in the selected contacts. The instantaneous PAF of the bistable EM and GM trials was further computed for each of the 10 contacts with the strongest alpha power, using similar methods as those for the EEG data analysis. The results showed that most of the selected contacts exhibited the trend of higher PAF in the bistable GM than bistable EM trials (Fig 4). For all the four patients, in the group mean of the ten contacts with maximal alpha amplitude, the PAF was significantly higher for the bistable GM percepts than the bistable EM percepts, using cluster-based permutation test (Fig 4A, patient 1: from about −500 to −110 ms relative to the presentation of Frame 1; Fig 4B, patient 2: from about −470 to −180 ms; Fig 4C, patient 3: from about −600 to −250 ms; and Fig 4D, patient 4: from about −410 to −300 ms, cluster-based correction, p < 0.05).

#### Prestimulus alpha frequency biased post-stimulus neural representation by inducing preactivation of the subsequently reported bistable percepts

We subsequently investigated how the PAF biases the neural representations of the spatially vs. temporally integrated percepts. The working hypothesis (Fig 1B) is that explicitly low PAFs will result in efficient perceptual inference on the EM percepts and inefficient perceptual inference on the GM percepts. On the other hand, explicitly high PAFs will result in efficient perceptual inference on the GM percepts and inefficient perceptual inference on the EM percepts. Accordingly, for the bistable GM trials with higher PAFs and the bistable EM trials with lower PAFs, we expected to observe biased neural representations of the predicted percepts in the neural signals shortly before the actual presentation of Frame 2 not only after, but also before the actual stimulus onset. To test the above hypothesis, we employed the temporal generalization method, a time-resolved decoding approach, to characterize how neural representations are dynamically transformed with the progress of time [52]. The classifiers used in the decoding analysis rely on boundaries through the high-dimensional activation space that maximally separate patterns of neural activity underlying different percepts (e.g. bistable EM vs. bistable GM). Classifier performance should be better if the two representations in the activation space are clearly separated [53] (Fig 5A).

**Fig 5.**
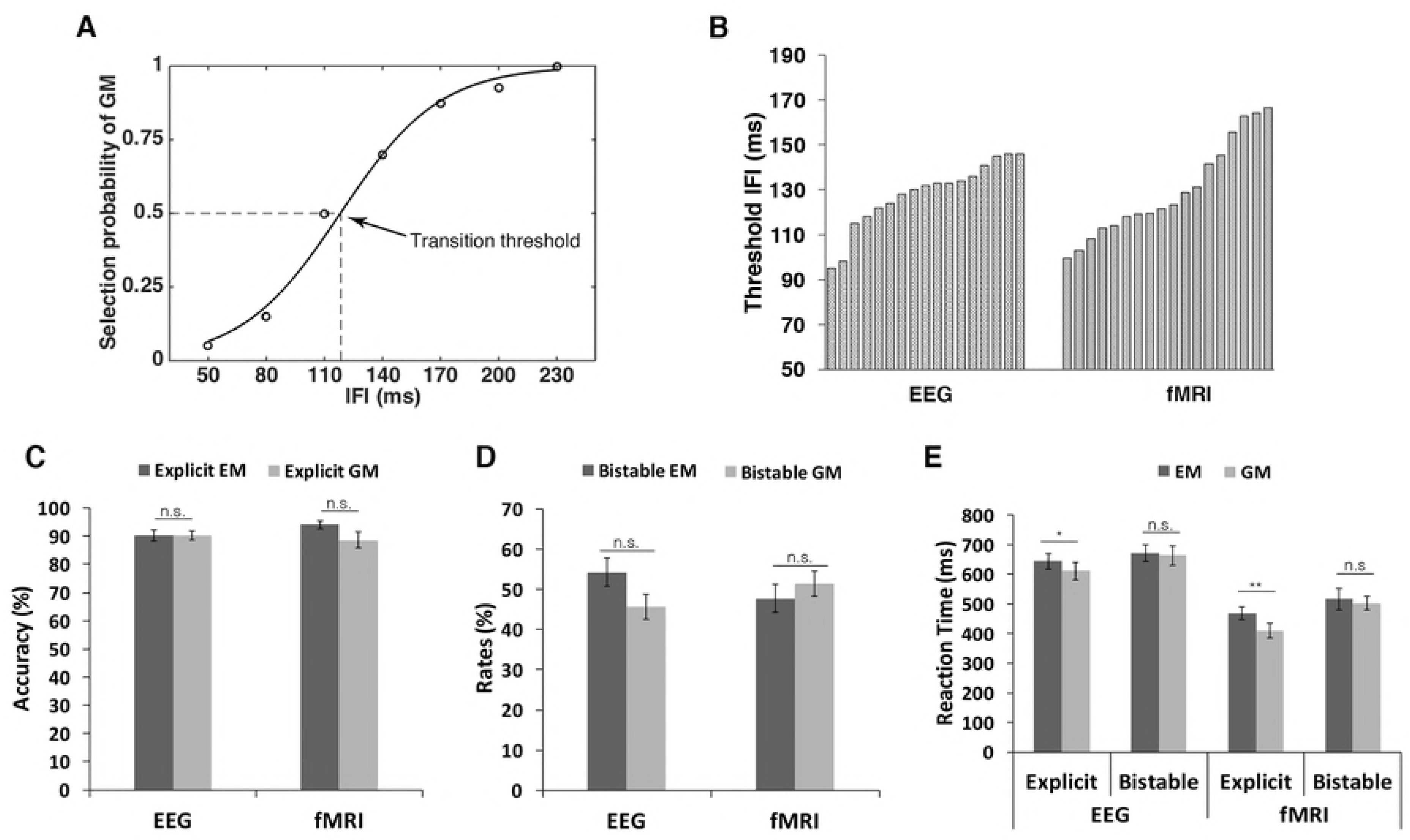
Decoding results of EEG data. (A) A hypothetical 2D activation space for the EEG signals representing the bistable EM and GM percepts. Activation patterns for each percept are projected onto a discriminant axis, which differentiates the two percepts. A decision boundary placed along the axis allows for classification between the bistable EM and GM percepts. The overlap of the Gaussian distributions reflects “decision noise”. More distant representations produce less noise, resulting in higher accuracy. (B) Selection of the bistable EM and GM trials according to the PAF. The high PAF GM trials (red dots in dark red shading) and the low PAF EM trials (blue dots in light blue shading) were designated as the efficient inference condition while the low PAF GM trials (red dots in light red shading) and the high PAF EM trials (blue dots in dark blue shading) were designated as the inefficient inference condition. (C) Temporal generalization matrices for the efficient inference condition. (D) Temporal generalization matrices for the inefficient inference condition. (E) Temporal generalization matrices for the differential contrast between the two conditions. Columns in the images are the time points the classifier was trained, and rows are the time points the classifier was tested. Color values indicate decoding accuracy in terms of AUC (C, D) or AUC difference (E). The Contour with the black line indicates the significant cluster, p < 0.05. (F) Decoding performance over time with training time around 50-100 ms after the onset of Frame 2. For the purpose of visualization, the figure shows a row in the temporal generalization matrix in (E) at the training time where we see a significant cluster of generalization difference before the onset of Frame 2. Significant generalization time points are indicated with horizontal black bars (cluster-based correction, p < 0.05). Time 0 indicates the onset of Frame 2. Shaded regions indicate ±1 SEM.

For the EEG data, all the bistable trials were first sorted according to the PAF, and then half split into the high PAF and the low PAF sessions. Subsequently, the bistable EM and GM trials in the high PAF session were selected as the high PAF EM and GM trials, respectively (Fig 5B). Similarly, the bistable EM and GM trials in the low PAF session were selected as the low PAF EM and GM trials, respectively (Fig 5B). To exclude the potential confounds caused by the different number of trials, the trial number in each of the above four types of trials was matched (see details in Materials and Methods). According to our hypothesis, the high PAF GM trials and the low PAF EM trials were designated as the efficient inference condition while the low PAF GM trials and the high PAF EM trials were designated as the inefficient inference condition (Fig 1B). For each condition, we calculated the temporal generalization matrix, which contained the decoding performance between the bistable EM and bistable GM trials over time (quantified by the area under the receiver operator characteristic, i.e. AUC, using the leave-one-out cross-validation method). If the process of perceptual integration simply depends on bottom-up sensory inputs, perceptual grouping between the two frames should happen only after the actual presentation of the second frame. We thus time-locked the temporal generalization matrix to the presentation of the second frame, to investigate how neural representations of the spatially vs. temporally integrated percepts were encoded with the progress of time relative to the presentation of Frame 2.

Neural representations of the bistable EM and GM percepts could be discriminated successfully after the presentation of the second frame for both the efficient inference condition (Fig 5C, from 10 ms to 380 ms after the onset of the second frame, cluster-based correction, p < 0.05) and the inefficient inference condition (Fig 5D, from 125 ms to 400 ms after the second frame, cluster-based correction, p < 0.05). By directly comparing the decoding performance between the two conditions, we found significantly higher decoding performance in the efficient than inefficient inference condition, shortly after the presentation of the second frame (the significant cluster from training time 10-170 ms and decoding time 0-170 ms, cluster-based correction, p < 0.05) (Fig 5E).

More interestingly, significant temporal generalization difference between the two conditions was revealed even before the presentation of the second frame (Fig 5E). In particular, the neural signals around 10-130 ms after the presentation of Frame 2 could be significantly better generalized to the neural signals 60-140 ms before the presentation of Frame 2 in the efficient than inefficient inference condition, and vice versa (140-160 ms before Frame 2 being generalized to 40-110 ms after Frame 2) (Fig 5E, cluster-based correction, p < 0.05). These generalized signals were further demonstrated when the decoder was trained around 50-100 ms after Frame 2 (Fig 5F): neural signals around 75-125 ms before Frame 2 were similar to those evoked by the actual onset of Frame 2, in the efficient inference condition. The above results together suggest that efficient perceptual inference based on the PAFs improved the readout of post-stimulus spatially vs. temporally integrated representations by pre-activating the percept-like signals even before the actual onset of Frame 2.

#### fMRI data: Prestimulus activity and network dynamics in the frontoparietal network predicted the outcome of bistable perceptual grouping

For the fMRI data, by directly contrasting the bistable EM and GM trials, we first identified the specific neural substrates underlying the bistable spatially vs. temporally integrated percepts. Compared to the Bistable GM trials, the Bistable EM trials induced stronger positive activations in the left intraparietal sulcus (IPS) (Fig 6A, upper panel, red; S2 Table), and stronger deactivations in the medial prefrontal cortex (MPFC) of the default-mode network (DMN) (Fig 6A, upper panel, blue; S2 Table). As shown in the mean parameter estimates extracted from the activated clusters (Fig 6A, lower panel), neural activity increased in the left IPS and decreased in the MPFC, specifically in the Bistable EM trials.

**Fig 6.**
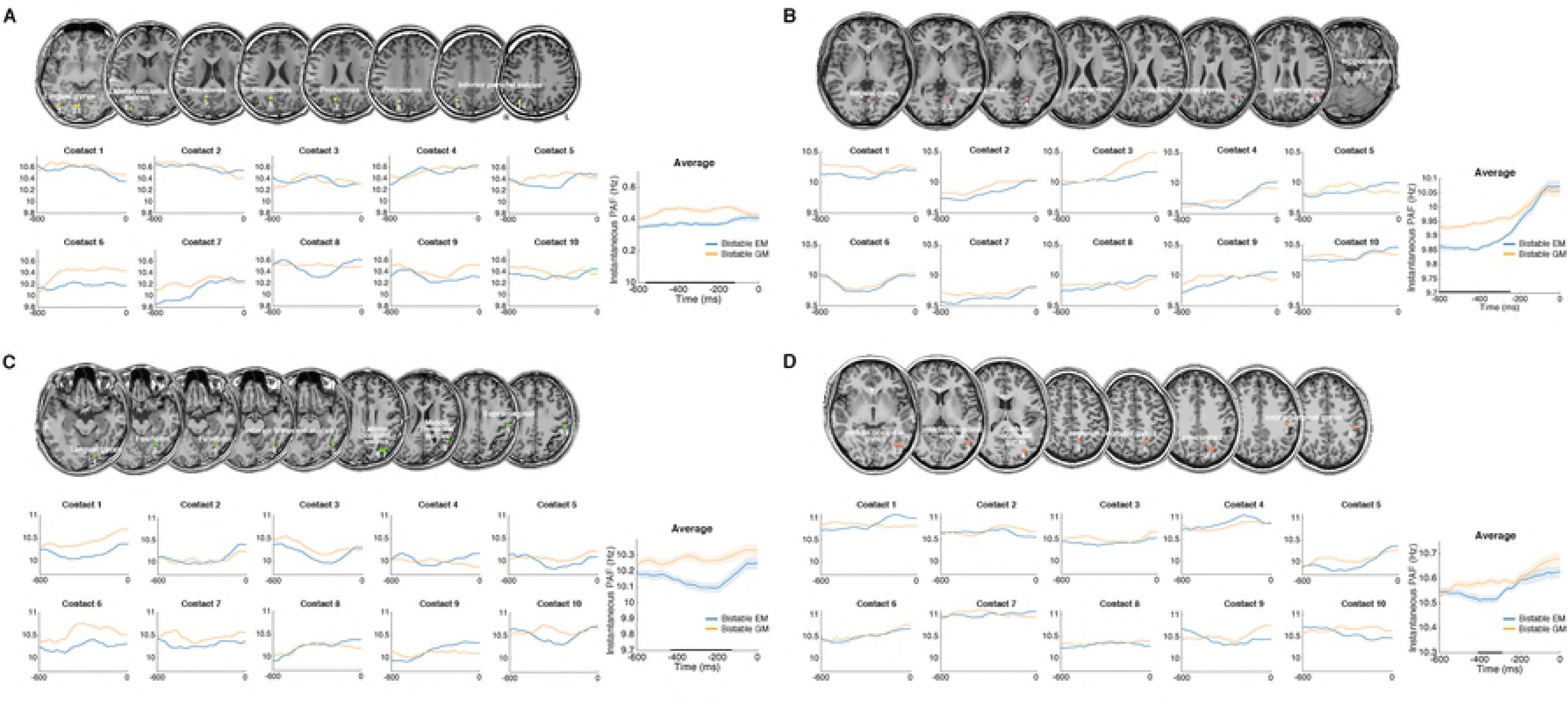
fMRI results: pre- and peri-stimulus neural activity and network dynamics in the bistable trials. (A) *Red*: left IPS showed significantly higher neural activity in the Bistable EM than Bistable GM trials. *Blue:* MPFC in default-mode network (DMN) was significantly more deactivated in the Bistable EM than Bistable GM trials. Parameter estimates in the four experimental conditions were extracted from the activated left IPS (*lower panel, left*) and the MPFC cluster (*lower panel, right*). The shaded condition is the Bistable EM condition that drives the significant neural contrasts. The error bars are ±1 SEM. (B) Increased prestimulus neural activity of the Bistable EM trials, compared to the prestimulus neural activity of the Bistable GM trials. Left IPS and bilateral IFG were significantly activated. Parameter estimates, indicating the height of neural activity in the previous trials (Trials N-1) of the bistable and the explicit trials, were extracted from the bilateral IFG. The height of neural activity in the trials prior to the Bistable EM trials was higher than in the trials prior to the Bistable GM trials (the two shaded conditions). (C) PPI analysis based on prestimulus neural activity in the left IPS, with the contrast “Bistable EM > Bistable GM” as the psychological factor. The left IPS showed enhanced neural coupling with sensorimotor and frontal areas during the prestimulus period of the Bistable EM trials, compared to the prestimulus period of the Bistable GM trials. For example, in a representative participant, mean corrected prestimulus neural activity in the supplementary motor area (SMA) is shown as a function of mean corrected prestimulus activity in the left IPS (i.e. the first principal component from a sphere of 4mm radius), for the Bistable EM trials (red dots and lines) and Bistable GM trials (blue dots and lines), respectively.

To further localize the specific neural regions in which prestimulus neural activity predicts the outcome of bistable perceptual grouping, we compared neural activity in the trials prior to the bistable EM and GM trials (see Materials and Methods). Left inferior parietal cortex and bilateral inferior frontal gyrus (IFG) showed significantly enhanced prestimulus neural activity of the bistable EM trials, compared to the bistable GM trials (Fig 6B, upper panel). For example, the extracted mean parameter estimates in bilateral IFG showed that prestimulus neural activity was higher for the bistable EM than the bistable GM trials, but was comparable between the explicit EM and the explicit GM trials (Fig 6B, lower panel; S2 Table). The left inferior parietal cortex showed similar patterns. The results suggested that enhanced neural activity in the frontoparietal network prior to the presentation of the bistable stimuli predicted the bistable EM percepts.

In addition to the height of prestimulus neural activity, prestimulus network dynamics in the frontoparietal attention network may play a critical role in predicting the outcome of bistable perceptual grouping as well. To address this, we compared the patterns of functional connectivity among the frontoparietal regions prior to the presentation of the bistable EM vs. GM trials. Since the left IPS exhibited specific selectivity towards the Bistable EM percepts during both pre- (Fig 6A) and post-stimulus (Fig 6B) period, we used the left IPS as the seed region to perform the network analysis, focusing on the prestimulus period.

Psychophysiological interaction (PPI) analysis treated prestimulus activity in the left IPS as the physiological factor, and the contrast between the two bistable percepts (Bistable EM vs. Bistable GM) as the psychological factor. In this way, we aimed to calculate how prestimulus changes in the functional connectivity of the left IPS predict the subsequent bistable EM vs. GM percepts. The results showed that prestimulus functional connectivity between the left IPS and the frontal regions was significantly enhanced for the bistable EM trials, compared to the bistable GM trials (Fig 6C; S3 Table). Therefore, the intrinsic organization status of the frontoparietal network prior to the presentation of the bistable stimuli predicted the subsequent subjectively perceived percepts.

## Discussion

By using EEG, intracranial recordings, and fMRI, we investigated how the frequency of alpha-band oscillations acts as the general neural dynamics that accommodate the temporal and spatial grouping during human perception, and more importantly how the brain makes predictions, based on intrinsic alpha frequency, to resolve perceptual ambiguity. At the behavioral level, comparable task performance/judgement difficulty was revealed between the bistable temporal and spatial grouping condition (Fig 2D, 2E). Therefore, any neuronal difference between the two bistable conditions cannot be attributed to differences in judgment difficulty. At the neural level, both within and between subjects, peak frequency of alpha oscillations in the occipitoparietal regions predicted the bistable temporal vs. spatial grouping (Fig 3 and 4). Moreover, efficient perceptual inference, based on spontaneous variance in PAF, induced a representation of the subsequently reported bistable percept in the neural signals shortly before the actual appearance of the second frame, indicating a preactivation of the subjectively perceived bistable percepts (Fig 5). In addition, variance in prestimulus activity and network dynamics in the frontoparietal network predicted the bistable temporal vs. spatial grouping, probably by modulating the frequency of the occipitoparietal alpha oscillations (Fig 6). Based on the above observations, we propose that the alpha frequency provides a general time window for perceptual grouping across both space and time, and perceptual inference based on intrinsic alpha frequency biased post-stimulus neural representations by inducing preactivation of the predicted percepts.

It has been proposed that perception is discrete and cyclic in a manner of perceptual cycles [15,16,41,54,55]. Accumulating recent evidence showed that perceptual performance depends on the frequency of the critical rhythm at around the onset time of stimuli [46,47,56]. A higher frequency of the brain oscillations should be equivalent to a faster frame rate of discrete perception, and vice versa. Accordingly, lower alpha frequency was reported to be associated with poorer temporal resolution [46,47], as if the slower frame rate of perception made two successive flashes more likely to fall within the same frame and thus be perceived as one [54]. Our results further delineate a general functional role of the alpha frequency in gating perceptual grouping across space and time. When the alpha frequency is relatively slow (i.e. longer alpha cycles) to cover both spatially and temporally segregated information segments with higher probabilities, temporal grouping dominates over spatial grouping, resulting in the EM percepts (Fig 1B, upper panel; Fig 3C, 3D). On the other hand, when the alpha frequency is relatively high (i.e. shorter alpha cycles) to cover only spatially segregated information segments with higher probabilities, spatial grouping dominates, resulting in the GM percepts (Fig 1B, lower panel; Fig 3C, 3D).

The intracranial data further confirmed the above alpha frequency effect in distributed visual areas including both the dorsal and ventral visual stream, such as the primary and secondary visual areas in the lingual gyrus, higher order areas in the fusiform gyrus, the lateral occipital cortex (LOC), the middle temporal gyrus (MT), and the intraparietal sulcus (Fig 4). The dorsal occipitoparietal areas, such as the inferior intraparietal sulcus, have been associated with perceptual integration of multiple elements and object representations [57,58]. The ventral visual areas, such as LOC, have been found to be involved in object recognition [59]. Moreover, it has been well documented that the MT area is highly responsive to visual motion and codes highly specialized representations of visual information [60–62], which is putative for generating apparent motion percepts [63,64]. The present results further suggest that the alpha frequency effect is a ubiquitous property of the visual system, which is involved in representing coherent object motion percepts. It has been revealed that neural oscillations could create temporal windows that favor the communication between neurons [65,66]. The common alpha frequency effect in distributed visual systems may drive the communication between neuronal groups in these areas to effectively encode and organize the dynamic visual inputs and induce coherent apparent motion percepts.

The generative models of perception, such as predictive coding [5,67–69], commonly consider the brain as an unconscious inference machine that uses hidden states to predict observed sensory inputs [68,70]. However, the physiological basis of the underlying computations remains unclear. In the present study, even with the sensory inputs (the two frames with a threshold IFI) being kept constant in the bistable condition, the subjective perception varies between the EM and GM percepts on a trial-by-trial base, thus suggesting fluctuations in prior predictions. The present Ternus display puts the brain under the explicit contextual information that there are two possible apparent motion percepts, i.e. GM vs. EM. Moreover, the alpha peak frequency, which provides a unified window for perceptual integration, is widely considered as one putative marker of an individual’s intrinsic state [71,72]. Therefore, perception is able to employ the current intrinsic alpha frequency to make predictions about the most possibly perceived apparent motion percepts (Fig 1B). Combining the specific prediction and forthcoming inputs, perceptual inference towards one specific percept will be made. Specifically speaking, since higher alpha frequencies make it more possible for the two frames to fall in different alpha cycles, higher PAFs will induce a more efficient inference towards the GM percepts and a less efficient inference towards the EM percepts (Fig 1B). On the other hand, since lower alpha frequencies make it more possible for the two frames to fall in the same alpha cycle, lower PAFs will induce a more efficient inference towards the EM percepts and a less efficient inference towards the GM percepts (Fig 1B). Indeed, our results showed that the peak alpha frequency not only casually predicted the outcome of bistable perceptual grouping (Fig 3C, 3D; Fig 4), but also modulated the fidelity of neural representations of the integrated percepts (Fig 5). Compared to the inefficient inference, neural representations of the bistable EM vs. GM percepts could be more robustly decoded immediately after the presentation of Frame 2 under the efficient inference (Fig 5C, 5D, and 5E), suggesting that efficient inference based on PAFs enhanced the fidelity of neural representations of the predicted percepts. More interestingly, under the efficient inference, the neural signals evoked by the actual presentation of the second frame could be readily read out from the neural signals even before the presentation of the second frame (Fig 5E, 5F), suggesting a preactivation of the predicted percepts. These results thus fundamentally advance our mechanistic understanding on how the alpha frequency builds up specific prediction signals for perceptual inference: perceptual predictions on the spatially vs. temporally integrated percepts are generated based on variations in the PAFs, which induces preactivated neural representations that resemble the neural representations evoked by the actual stimuli.

In addition to the PAF effect, the fMRI results showed that both increased prestimulus neural activity (Fig 6B) and increased prestimulus network dynamics (Fig 6C) in the frontoparietal network predicted the subsequent bistable EM (temporal grouping), rather than GM (spatial grouping), percepts. Moreover, the enhanced frontoparietal activations and DMN deactivations during the bistable EM trials (Fig 6A) indicated that bistable temporal grouping was more attention demanding than bistable spatial grouping [73,74], probably because more information segments fall in the same alpha cycle during temporal than spatial grouping. Together with the EEG data, the present results thus suggested that the bistable temporal grouping is associated with enhanced attention in the frontoparietal network (Fig 6) and decreased posterior occipitoparietal PAFs (Fig 3D, Fig 4). The peak alpha frequency has been considered as a predictor of attention [75]: slower alpha oscillations reflect attentional processes. It has been well documented that the frontoparietal attention network generates and maintains a top-down attention signal to selectively bias posterior occipital alpha activity [76,77]. A recent MEG study revealed that visuospatial attention was associated with the long-range alpha synchronization among frontal, parietal and occipital visual regions [78], and a simultaneous EEG-fMRI study revealed that the long-range alpha synchrony was intrinsically linked to activity in the frontoparietal attentional control network [79]. Moreover, alpha lateralization was disrupted after processing in the IPS was selectively interrupted by repetitive transcranial magnetic stimulation (rTMS) [80], which provides causal evidence on the functional link between activity in the frontoparietal network and modulation of posterior alpha activity. Therefore, when visual perception relies on higher order areas in the frontoparietal attentional control network as in the present bistable temporal grouping condition (Fig 6), a relatively slower occipitoparietal alpha rhythm (Fig 3D, Fig 4) would emerge as a result of the top-down inter-areal modulation from the frontoparietal regions [81].

To summarize, by creating a bistable/ambiguous display in which subjective perception fluctuates between temporally vs. spatially integrated percepts, we showed that the occipitoparietal alpha frequency defines a unified temporal window for perceptual grouping across space and time. Moreover, efficient perceptual inference, based on the alpha frequency, induces preactivated neural representations of the predicted percepts already before the actual presentation of the critical stimuli.

Therefore, perceptual inference employs PAF-induced predictions to resolve perceptual ambiguity.

## Materials and Methods

### Participants

Nineteen participants (12 females, mean age of 19.6 years old) took part in the EEG experiment. Another group of twenty participants (12 females, mean age of 23.4 years old) took part in the fMRI experiment. Two participants in the EEG experiment were discarded due to excessive eye movement artifacts. One participant in the fMRI experiment was discarded due to low accuracy (less than 70%) in the Explicit conditions, and another participant was discarded due to the excessive head movements during the scanning. Therefore, 17 participants in the EEG experiment and 18 participants in the fMRI experiment were included for further analysis.

Additionally, four patients (two males, mean age of 24 years old) undergoing intracranial recordings with stereotactically implanted multi-lead electrodes (Guangdong Sanjiu Brain Hospital, China) for epilepsy treatment participated in the present study. The placement of the depth electrodes was based solely on the clinical needs for the treatment of the patients, and was thus independent of the purpose of the present study. Although anatomical locations of the electrodes were different in each patient, we included the patients whose electrodes were implanted in the occipital and parietal regions. Patients who had destructive lesions such as tumor or encephalomalacia were excluded. This study did not add any invasive procedure to the intracranial recordings. All the participants gave their informed consent prior to the experiment in accordance with the Declaration of Helsinki. All of them were right-handed, with normal or corrected-to-normal visual acuity. The fMRI, the EEG, and the patient experiments were all approved by the Ethics Committee of School of Psychology, South China Normal University.

### Stimuli

Visual stimuli consisted of two consecutively presented frames of stimuli (Frame 1 and Frame 2), and each frame was presented for 30 ms (Fig 1). There was a blank period between the two frames, i.e. the inter-frame interval (IFI). The IFI could be either explicitly short at 50 ms, or explicitly long at 230 ms, or at the transition threshold. Each frame contained two horizontally arranged black discs (1.6° of visual angle in diameter) on a gray background. The center-to-center spatial distance between the two discs was 3° of visual angle. The two frames shared one common disc location at the center of the display. The location of the lateral disc of the first frame, either on the left or the right side of the shared central disc, was always opposite to the lateral disc of the second frame (Fig 1). Specifically speaking, Frame 1 with left and central discs and Frame 2 with right and central discs induced rightward apparent motion; Frame 1 with right and central discs and Frame 2 with left and central discs induced leftward apparent motion. The same set of stimulus parameters was adopted for the fMRI, the EEG, and the patient experiments. Depending on the IFI and participants’ online judgements in the bistable trials, there were four types of experimental trials: (1) the explicit EM trials (“Explicit EM”) with the short IFI of 50 ms; (2) the explicit GM trials (“Explicit GM”), with the long IFI of 230 ms; (3) the bistable trials with the threshold IFI, which were judged by the participants as the EM trials (“Bistable EM”); and (4) the bistable trials with the threshold IFI, which were judged by the participants as the GM trials (“Bistable GM”).

### Psychophysical Procedures

To specify the 50% threshold of IFI for the bistable condition for each individual subject, we asked each participant to perform a psychophysical pretest before the main experiment. Prior to the psychophysics test, participants were shown demos of the explicit EM and GM conditions, and performed a practice block with only explicit EM and GM trials until the accuracy reached no less than 95%. During the formal psychophysics test, the first frame was presented for 30 ms. After a variable IFI (seven levels: 50, 80, 110, 140, 170, 200, or 230 ms), the second frame was presented for 30 ms as well. For each IFI condition, the percentage of GM reports was collapsed over the leftward and rightward motion directions. The seven data points (one for each IFI) were fitted into a psychometric curve using a logistic function [82]. The transition IFI threshold, i.e. the point at which EM and GM were reported with equal possibility, was calculated by estimating the 50% performance point on the fitted logistic function for each participant [82]. The individual transition threshold derived from the psychophysics test was then used as the IFI in the bistable trials of the subsequent main experiment. Different from the EEG and fMRI experiment, in the intracranial experiment, an adaptive staircase procedure [83] was adopted to find the individual IFI threshold at which 50% of the stimuli were perceived as GM.

### Main Experiment Procedures

Participants were instructed to fixate at a central fixation throughout the experiment without moving their eyes. The experimental task was to discriminate the two types of motion by pressing two pre-specified buttons on the response pad using the thumb of each hand, respectively. The mapping between the two response buttons and the two types of apparent motion percept was counterbalanced between participants.

In each trial, the first frame was presented for 30 ms, and after a variable IFI (50 ms, 230 ms, or the individual IFI threshold), the second frame was presented for another 30 ms. The fMRI experiment consisted of 440 trials in total, including 80 explicit EM trials, 80 explicit GM trials, 160 bistable trials, and 120 null trials. The null trials, in which only the central fixation cross was presented, were used as the implicit baseline. The participants were asked to rest for a short period of time [11 s, i.e. five repetition times (TRs)] after every 6 minutes’ task performance, which made three short periods of rest in total. During the three short rest periods, the scanner kept running and a visual instruction “rest” was presented on the center of the screen throughout. One TR after the disappearance of the “rest” instruction, the behavioural task resumed. The EEG experiment consisted of four blocks, and each block included 40 explicit EM trials, 40 explicit GM trials, and 80 bistable trials, which were intermixed randomly, resulting in 640 experimental trials in total. A rest break was allowed between blocks. For the fMRI and EEG experiment, each trial was followed by a time interval that was selected randomly among 2000, 2250, 2500, 2750, and 3000 ms. In the intracranial experiment, there were four blocks of 80 trials (320 trials in total), 10% of which were explicit EM and GM trials. The inter-trial interval varied randomly between 1.5 and 2.5 s. In all the three experiments, the temporal order of all the trials was randomized for each participant individually to avoid potential problems of unbalanced transition probabilities. All participants completed a training section of 5 min before the recording.

### Recording and Preprocessing of the EEG Data

EEGs were continuously recorded from 64 Ag/AgCl electrodes (10–20 System) with BrainAmp DC amplifiers (low-pass = 100 Hz, high-pass = 0.01 Hz, and sampling frequency = 500 Hz). The vertical electro-oculogram was recorded by one electrode under the participants’ left eyes. All the electrode impedances were kept below 5kΩ.

Signals were referenced online to the unilateral mastoid. Offline processing and analysis were performed using EEGLAB [84] and customized scripts in MATLAB (MathWorks). Data were down sampled to 160 Hz, re-referenced to the average reference, epoched from −800 ms before the first frame to 1000 ms after the first frame for the subsequent alpha frequency analysis, and re-epoched from −500 ms to 500 ms relative to the presentation of the second frame for the decoding analysis. Trials containing visually identified eye movements or muscle artifacts were excluded manually. Visually identified noisy electrodes were spherically interpolated.

### Acquisition and Preprocessing of the Intracranial Data

Ten to thirteen semirigid, multi-lead electrodes were stereotactically implanted in the four participants, respectively. All the electrodes have a diameter of 0.8 mm and contain 10-16 2mm-wide and 1.5mm-apart contacts. The precise anatomical location of each contact was identified by co-registering each participant’s post-implantation CT with the pre-implantation 3D T1 image, using rigid affine transformations derived from FSL’s FLIRT algorithm [85]. Intracranial recordings were conducted using commercial video-intracranial monitoring system. The data were bandpass-filtered online from 0.1 to 300 Hz and sampled at 1000 Hz, using a reference contact located in the white matter. For the offline analysis, recording signals were down sampled to 500 Hz. Contacts in the epileptogenic zones were excluded from further analyses.

Each contact was re-referenced with respect to its direct neighbor, i.e. bipolar montage, to achieve high local specificity by removing effects of distant sources that spread equally to adjacent sites through volume conduction. All the data were epoched from −800 to 1000 ms relative to the presentation of the first frame.

### Acquisition and Preprocessing of the fMRI Data

A Siemens 3T Trio system with a standard head coil at Beijing MRI Center for Brain Research was utilized to obtain T2*-weighted echo-planar images (EPIs) with blood oxygenation level-dependent contrast. The matrix size was 64 x 64 mm^3^ and the voxel size was 3.4 x 3.4 x 3 mm^3^. Thirty-six transversal slices of 3 mm thickness that covered the whole brain were acquired sequentially with a 0.75 mm gap (TR = 2.2 s, TE = 30 ms, FOV = 220 mm, flip angle = 90°). There was a single run of functional scanning, including 524 EPI volumes. The first five volumes were discarded to allow for T1 equilibration effects.

Data were preprocessed with Statistical Parametric Mapping software SPM12 (Wellcome Department of Imaging Neuroscience, London, http://www.fil.ion.ucl.ac.uk). Images were realigned to the first volume to correct for inter-scan head movements. The mean EPI image of each participant was then computed and spatially normalized to the MNI single participant template using the “unified segmentation” function in SPM12. This algorithm is built on a probabilistic framework that enables image registration, tissue classification, and bias correction to be combined within the same generative model. The resulting parameters of a discrete cosine transform, which define the deformation field necessary to move individual data into the space of the MNI tissue probability maps, were then combined with the deformation field transforming between the latter and the MNI single participant template. The ensuing deformation was subsequently applied to individual EPI volumes. All images were thus transformed into standard MNI space and resampled to 2 x 2 x 2 mm^3^ voxel size. The data were then smoothed with a Gaussian kernel of 8 mm full-width half-maximum to accommodate inter-participant anatomical variability. Data were high-pass-filtered at 1/128 Hz and analyzed with a general linear model as implemented in SPM12. Temporal autocorrelation was modeled using an AR (1) process.

### Analysis of the Behavioral Data

For the behavioral data in the EEG, intracranial and fMRI experiment, omissions and trials with reaction times (RTs) 3 standard deviations (SDs) away from the mean RT in each condition were first excluded from further analysis. For both the EEG and fMRI experiment, paired t-tests were performed to test the difference in the accuracy rates between the two types of explicit trials, the proportions of EM and GM trials in the bistable condition, and the mean reaction times (RTs) for the two explicit and the two bistable conditions, respectively.

### Alpha Frequency Analysis of the EEG Data

For all the electrodes in all the participants, a power spectrum (from 5 Hz to 30 Hz) was obtained through a Fast Fourier Transform (FFT) of all the trials (from −800 to 0 ms relative to the presentation of the first frame). An amplitude topographic map of the most prominent frequency band in the power spectrum was obtained. For each participant, the individual peak alpha frequency was determined as the value corresponding to the maximum peak frequency from the 800 ms of data prior to the presentation of the first frame within the 8–13 Hz range for the selected posterior electrodes. The Pearson product-moment correlation between the width of individual alpha cycle and the individual IFI threshold obtained from the psychophysical procedures was then calculated.

Instantaneous PAF was analyzed using the methods and code developed by Cohen [51]. First, individual data from the electrodes showing the strongest alpha amplitude were epoched from −800 to 0 ms relative to the presentation of the first frame. To avoid edge artifacts at the stimulus onset due to filtering, data from each trial were copied and flipped from left to right and appended to the right side of the original data. Epochs were filtered between 8 and 13 Hz with a zero-phase, plateau-shaped band-pass filter with 15% transition zones. Phase angle time series were extracted from the filtered data with a Hilbert transform. The temporal derivative of the phase angle time series describes how phase changes over time, and thus corresponds to the instantaneous frequency in Hz (when scaled by the sampling rate and 2 *π*). Since noises in the phase angle time series can cause sharp, non-physiological responses in the derivative, the instantaneous frequency was filtered with a median filter with an order of ten and a maximum window size of 400 ms: data were median filtered ten times with ten time windows ranging from 10 to 400 ms prior to averaging across trials. Since this analysis considers changes only in the instantaneous phase of the data, it is mathematically independent from the amplitude of the oscillation, except where amplitude is equal to zero and phase is undefined. Subsequently, the instantaneous PAFs were averaged across bistable EM and GM trials, respectively.

### Decoding Analysis of the EEG Data

Multivariate decoding techniques were further adopted to investigate how the PAF affects the representation contents of the bistable EM and GM percepts with the progress of time. We first calculated the PAF for each bistable trial by averaging the calculated instantaneous alpha frequency across the time points which showed a significant difference between the bistable EM and GM precepts at the group-level (−550 ∼ −210 ms relative to the presentation of Frame 1; see Fig 3D). Subsequently, all the bistable trials are sorted according to the PAF, and half split into the high PAF and the low PAF trial sessions. The bistable GM trials in the high PAF session and the bistable EM trials in the low PAF session were selected as the two types of trials in the efficient inference condition; the bistable EM trials in the high PAF session and the bistable GM trials in the low PAF session were selected as the two types of trials in the inefficient inference condition. To exclude potential confounds caused by different number of trials upon comparing different conditions, we matched the trial count in the above four types of trials, by randomly selecting a sub-sample of trials from the conditions with more trials. We then applied a multivariate linear discriminant analysis to characterize the temporal dynamics that discriminated between the subjectively perceived bistable EM vs. GM percepts for the efficient and inefficient inference condition, respectively.

Classifications were based on the regularized linear discriminant analysis to identify a projection in the multidimensional EEG data, *x*, that maximally discriminated between the two representations across all stimulus levels. Each projection is defined by a weight vector, *w*, which describes a one-dimensional projection *y* of the EEG data y = ∑_*i*_*w_i_x_i_* + *c*, with i summing over all channels and c a constant. Theregularization parameter was optimized in preliminary tests and kept fixed for all the analyses. The decoding analysis was performed in a time-resolved manner by applying to each time point sequentially, resulting in an array of classifiers, e.g. *w*(t1), *w*(t2), *w*(t3) and so on. To improve the signal-to-noise ratio, the data were first averaged within a time window of 50 ms centered around the time point of interest. Subsequently, the classifier performance was assessed not only at the time point used for training, e.g. classifier *w*(t1) was tested at t1, *w*(t2) was tested at t2, and so on, but also on data from all the other time points, e.g. classifier *w*(t1) was tested on all the time points t1, t2, t3, and so on. The performance of the classifier was quantified using the receiver operator characteristic (ROC), based on leave-one-out cross-validation within each participant. The above procedure resulted in a (training time) × (decoding time) temporal generalization matrix per condition. To further investigate how the discriminability between the two bistable percepts differentially generalized across time between the efficient and inefficient inference condition, we compared the temporal generalization matrix between the efficient and inefficient inference condition, by subtracting one from the other to obtain a difference matrix.

### Alpha Frequency Analysis of the Intracranial Data

We first extracted the averaged alpha amplitude during the prestimulus period (−800 to 0 ms relative to the first frame) for each contact in the same manner as for the EEG analysis (using FFT). The first 10 contacts with the highest alpha amplitude (8–13 Hz) were then selected as regions of interest for each patient. Subsequently, we adopted similar methods and procedures as in the EEG analysis to calculate the prestimulus instantaneous frequency for the bistable EM and GM trials for each contact, which was subsequently averaged across the ten contacts.

### Statistical Testing of the Neurophysiology Data

The difference between two conditions was statistically tested using nonparametric cluster-based permutation tests, which were implemented in customized scripts in MATLAB (MathWorks). Specifically speaking, paired t-tests were first calculated for the two conditions. Elements that passed a threshold value corresponding to a p value of 0.05 were marked, and neighboring marked elements were identified as clusters. Cluster-based correction was applied when multiple time points were tested (Fig 3D; Fig 4; Fig 5C-F): data were randomly shuffled 1,000 times (500 times in the decoding analysis); and for each shuffle, the largest cluster size was entered into a distribution of cluster sizes, which was expected under the null hypothesis. Clusters in the real data were considered as statistically significant only if they exceeded the size of 95^th^ percentile of the null distribution of clusters, at α = 0.05.

### Statistical Analysis of the fMRI Data

At the individual level, the general linear model (GLM) was used to construct a multiple regression design matrix. The four experimental conditions were modeled as regressors of interest: Explicit EM, Explicit GM, Bistable EM, and Bistable GM. The four types of event were time locked to the onset of the first frame in each trial by a canonical synthetic hemodynamic response function and its first-order time derivative with an event duration of 0 s. In addition, all the omission trials and the outlier trials in which RTs were outside of the mean RT ± 3SD were modeled separately as another regressor. The six head movement parameters derived from the realignment procedure were also included as confounds. Parameter estimates were subsequently calculated for each voxel using weighted least-square analysis to provide maximum likelihood estimators based on the temporal autocorrelation of the data. No global scaling was applied.

For each participant, simple main effects for each of the four experimental conditions were computed by applying appropriate “1 0” baseline contrasts, that is, experimental conditions vs. implicit baseline (null trials). The four first-level individual contrast images were then fed into a within-participants ANOVA at the second group level employing a random-effects model (flexible factorial design in SPM12 including an additional factor modeling the subject means). In the modeling of variance components, we allowed for violations of sphericity by modeling non-independence across parameter estimates from the same subject and allowing unequal variances between both conditions and participants using the standard implementation in SPM12. We were particularly interested in the differential neural activity between the two types of bistable trials (Bistable EM vs. Bistable GM). Areas of activation were identified as significant only if they passed a conservative threshold of p < 0.005, family-wise error (FWE) corrected for multiple comparisons at the cluster level, with an underlying voxel level of p < 0.005, uncorrected [86].

### Statistical Analysis on Prestimulus Neural Activity of the fMRI Data

To investigate how the prestimulus neural activity predicted the outcome of bistable perceptual grouping, a new GLM model was estimated. Given that the ITI was jittered between 2000-3000 ms and one third of all the trials were null trials, the prestimulus periods of all the experimental trials were long enough and adequately jittered for the present statistical analysis on prestimulus neural activity. In the new GLM model, four types of new events were time locked to the time points after the participants made their responses in the preceding trials (‘Trials N-1’) of the four types of experimental trials, i.e. the prestimulus preparation period of the current trial (‘Trials N’). All the outliers, errors, and missed trials, and trials preceded by outliers and errors were separately modeled as another regressor. In this way, parameter estimates in each of the four newly defined critical neural events indicate the height of prestimulus preparation neural activity prior to the actual presentation of the explicit EM, the explicit GM, the bistable EM, and the bistable GM stimuli. Brain regions of activation were identified as significant only if they passed a conservative threshold of p < 0.005 family-wise error (FWE) correction for multiple comparisons at the cluster level, with an underlying voxel level of p < 0.005, uncorrected.

### Psychophysiological Interaction (PPI) Analysis on Prestimulus Neural Activity of the fMRI Data

For the PPI analysis, prestimulus neural activity (time locked to the responses in ‘Trials N-1’) in the left IPS was used as the physiological factor, and the contrast of “Bistable EM vs. Bistable GM” as the psychological factor. For each participant, the neural contrast of “Bistable EM vs. Bistable GM” was first calculated in the individual level GLM. Subsequently, each participant’s individual peak voxel in the left IPS was determined as the maximally activated voxel within a sphere of 16mm radius (i.e., twice smoothing kernel) around the coordinates of the peak voxel from the second-level group analysis (Fig 6A). Individual peak voxels from every participant are located in the same anatomical structure (left IPS MNI coordinates: x = −33 ± 6, y = −37 ± 7, z = 42 ± 6). Next, the left IPS time series in every participant were extracted from a sphere of 4 mm radius around the individual peak voxels. The PPI term was created for each participant by multiplying the deconvolved and mean-corrected BOLD signal in the given ROI (i.e. the physiological variable) with the psychological variable of interest (i.e. “Bistable EM vs. Bistable GM”). After convolution with the HRF, mean correction, and orthogonalization, three regressors (the PPI term, the physiological variable, and the psychological variable) were entered into the GLM to reveal areas in which neural activations were predicted by the PPI term, with the physiological and the psychological regressors being treated as confound variables. The PPI analysis was first carried out for each participant, and then entered into a random-effects group analysis. Statistical significance was set to p < 0.005, uncorrected at the voxel level, with the cluster extent exceeding 100 voxels.

## Acknowledgments

We thank Klaartje Heinen for her useful comments. Q.C. is supported by the Program for New Century Excellent Talents in the University of China (NCET-12-0645) and Changjiang Scholars Program (2016). L.S. is supported by China Scholarship Council and the Top-Notch Graduate Foundation of South China Normal University. B.H. is supported by Fyssen Foundation. L.C. is supported by Grants from the Natural Science Foundation of China (61621136008, 61527804) and DFG TRR-169 in project Crossmodal Learning.

## Supporting Information

**S1 Table. Patient information and behavioral performance in the intracranial experiment.** Following information is reported for each patient: the epileptogenic zone (EZ) identified by the clinical investigation; the number of implanted electrode shafts (EL); the total number of electrode contacts (CH). Task performance: accuracy rates (AR) of the Explicit EM and GM condition; reported rates of GM in the Bistable condition.

**S2 Table. Brain activations in the main Fig 6A, B.** Brain regions showing significant relative increases of BOLD response associated with the Bistable EM and Bistable GM trials before and after the presentation of the first frame.

**S3 Table. Brain activations in the main Fig 6C.** Brain regions that showed higher pre-stimulus functional connectivity with the left IPS (−32, −38, 38) in the Bistable EM than Bistable GM trials.

S1 Video. Demo of explicit EM.

S2 Video. Demo of explicit GM.

